# Fine-scale seascape genomics of an exploited marine species, the common cockle *Cerastoderma edule*, using a multi-modelling approach

**DOI:** 10.1101/724062

**Authors:** Ilaria Coscia, Sophie B. Wilmes, Joseph E. Ironside, Alice Goward-Brown, Enda O’Dea, Shelagh K. Malham, Allan D. McDevitt, Peter E. Robins

## Abstract

Population dynamics of marine species that are sessile as adults are driven by oceanographic dispersal of larvae from spawning to nursery grounds. This is mediated by life-history traits such as the timing and frequency of spawning, larval behaviour and duration, and settlement success. Here, we use 1725 single nucleotide polymorphisms (SNPs) to study the fine scale spatial genetic structure in the commercially important cockle species *Cerastoderma edule* and compare it to environmental variables and current-mediated larval dispersal within a modelling framework. Hydrodynamic modelling employing the NEMO Atlantic Margin Model (AMM15) was used to simulate larval transport and estimate connectivity between populations during spawning months (April - September), factoring in larval duration and seasonal variability of ocean currents. Results at neutral loci reveal the existence of three separate genetic clusters (mean *F*_ST_=0.021) within a relatively fine spatial scale in the northwest Atlantic. Environmental association (EA) analysis indicates that oceanographic currents and geographical distance between the populations explain over 20% of the variance observed at neutral loci, while genetic variance (71%) at outlier loci was explained by sea surface temperatures extremes. These results fill an important knowledge gap in the management of a commercially important, overexploited species, and bring us closer to understanding the role of larval dispersal in connecting populations at a fine geographical scale.

## 1. Introduction

The assumption that genetic homogeneity predominates in marine organisms due to the lack of physical barriers and high dispersal potential at all life stages has been challenged in recent years (Allendorf, 2017). The advent of genomic markers generated by scanning the whole genome of an organism has equipped researchers with the necessary statistical power to detect more fine-scale population differentiation within the marine realm (Benestan et al., 2015; Maroso et al., 2018). Furthermore, genome scans have allowed a shift in focus from neutral variation to local adaptation, which is one of the main drivers of structure within marine populations (Araneda, Larraín, Hecht, & Narum, 2016; Nielsen et al., 2012; Woodings et al., 2018) and indicates the potential for resilience to future environmental change (Razgour et al., 2019). Connectivity between marine populations is central to their health and resilience to external pressures such as parasites and pathogens (Rowley et al., 2014), pollution, human exploitation and climate change over ecological and evolutionary timescales (Burgess, Bowring, & Shen, 2014; Cowen & Sponaugle, 2009; Gimenez, 2019). For these reasons, it is vital to distinguish between neutral variation and local adaptation when attempting to understand what drives the observed population structure in marine organisms.

A seascape genomics approach is particularly valuable in this context (Selkoe et al., 2016). Genetic and genomic data can be used in conjunction with environmental variables such as sea water temperature, salinity, water depth, irradiance, turbidity, and sediment type (Viricel & Rosel, 2014) against the neutral differentiation determined by the equilibrium between larval dispersal and effective population size (Coscia, Robins, Porter, Malham, & Ironside, 2013; Young et al., 2015). This can provide new insights for interpreting genetic patchiness in relation to specific environmental features (Benestan et al., 2016; Bernatchez et al., 2019; Truelove et al., 2017).

In particular, seascape genomics has proven to be extremely useful when considering exploited species, as it has the potential to inform management and aid sustainable exploitation by enabling management units to be defined (Bernatchez et al., 2017; Teacher, André, Jonsson, & Merilä, 2013). Despite intense exploitation of many marine species, their management rarely takes into account genomic information (Bernatchez et al., 2017; ICES, 2018) and exploited aquatic invertebrates (shellfish) in particular receive little attention from policy makers and stakeholders in comparison with fish (Elliott & Holden, 2017). In the Irish Sea, shellfish represent 80% of the total landings per year, with an economic value of £46.6 M (Elliott & Holden, 2017). Among these, the common cockle *Cerastoderma edule* fisheries are some of the most valuable fisheries for both Ireland and the United Kingdom (Dare, Bell, Walker, & Bannister, 2003; Hervas, Tully, Hickey, O’Keeffe, & Kelly, 2007) and are of high socio-economic importance, valued at £3.3 M for Wales alone (Elliott & Holden, 2017).

The common cockle has both ecological and commercial importance, providing an important food source for wading birds in addition to employment for coastal communities (Flach & de Bruin, 1994; Hickin, 2019). *C. edule* occurs in intertidal soft sediment regions of the eastern Atlantic, from Norway to Senegal. It can live for six to ten years and is characterized by high fecundity and high dispersal potential due to a pelagic larval phase which lasts for 24 to 35 days following spawning from May to August (Malham, Hutchinson, & Longshaw, 2012). In recent decades, cockle stocks have declined across Europe, with production falling from 108,000 tons in 1987 to less than 25,000 tons in 2008 (Martínez, Mendez, Insua, Arias-Pérez, & Freire, 2013). Cockle declines have been attributed to different factors in different locations, such as overharvesting (Wolff, 2005) and parasitic infections (Longshaw & Malham, 2015; Thieltges, 2006). In the UK, recurrent mass mortalities have occurred at several long-established cockle fisheries, resulting in significant economic losses (Woolmer, 2013). These mortalities have not been attributed to any single environmental factor and interactions between multiple stress factors are suspected (Callaway, Burdon, Deasey, Mazik, & Elliott, 2013; Malham et al., 2012). Sustainable management of the common cockle is hindered by poor understanding of their population connectivity. Analysis of microsatellite and mitochondrial DNA markers suggests weak barriers to gene flow between *C. edule* populations along the North European coast (Coscia et al., 2013; Martínez, Freire, Arias-Pérez, Mendez, & Insua, 2015). However, these markers lack sufficient resolution to investigate connectivity at the finer scales relevant to fisheries management (Bernatchez et al., 2017).

Given the major logistical challenges of directly quantifying larval connectivity, efforts have focused on simulating ocean hydrodynamics to deduce larval dispersal probabilities (Cowen, Gawarkiewicz, Pineda, Thorrold, & Werner, 2007; Paris, Cowen, Claro, & Lindeman, 2005; Robins, Neill, Giménez, Jenkins, & Malham, 2013). This approach identifies well-connected population groups as well as weakly-connected, partially-connected or isolated populations. These simulations highlight the importance of local and mesoscale hydrodynamics interacting with species-specific larval behaviours in driving population persistence (Bode et al., 2019; Botsford et al., 2009; North et al., 2008; Robins et al., 2013) and recovery from stock decline (Gimenez, 2019), indicating that the capacity of a population to recover from mass mortalities is contingent on the scale of disturbance relative to the scale of connectivity. Usual circulation patterns and, hence connectivity, can be modulated by severe wind and wave conditions. Previous larval dispersal studies have predicted that given atypical meteorological conditions during spawning events, new connectivity routes can be established (e.g. by reversing the Celtic Sea front circulation; Hartnett, Berry, Tully, & Dabrowski, 2007), or by large distance displacements from expected routes affecting sea turtles: (Monzón-Argüello C. et al., 2012).

In the present study, a seascape genomics approach using single nucleotide polymorphisms (SNPs) is employed, for the first time, to resolve patterns of population structure of the common cockle between estuaries within a commercially active area (the Irish and Celtic Seas). An additional aim is to investigate the influence of environmental factors, including oceanic currents and water temperature upon the cockle’s population structure. To do this, larval transport between sites is estimated and connectivity matrices are derived from oceanographic modelling, accounting for variability due to bio-physical parameters, i.e. spawning date and larval duration. Finally, Environmental Association analysis (EA) is used in order to understand the interplay between larval dispersal and abiotic factors in shaping the population connectivity measured with genomic markers.

## 2. Materials and methods

### 2.1. Sampling and DNA extraction

Cockles were collected between 2010 and 2011 from seven locations off the coasts of Ireland (Bannow Bay [BAN] and Flaxfort strand [FLX]) and Britain (Burry Inlet [BUR], Gann Estuary [GAN], Dyfi estuary [DYF], Red Wharf Bay [RWB] and Dee estuary [DEE]) (Fig. 1 and Table 1). Genomic DNA was extracted from ethanol-preserved and frozen tissue using the DNeasy Blood and Tissue Kit with an additional RNaseA step (QiagenⒸ), as per the manufacturer’s protocols. The quality and quantity of the extracted DNA was assessed by gel electrophoresis (1% agarose) as well as Qubit dsDNA HS (high sensitivity, 0.2 to 100 ng) Assay Kit on a Qubit 3.0 fluorometer according to the manufacturer’s protocols.

**Figure 1:**
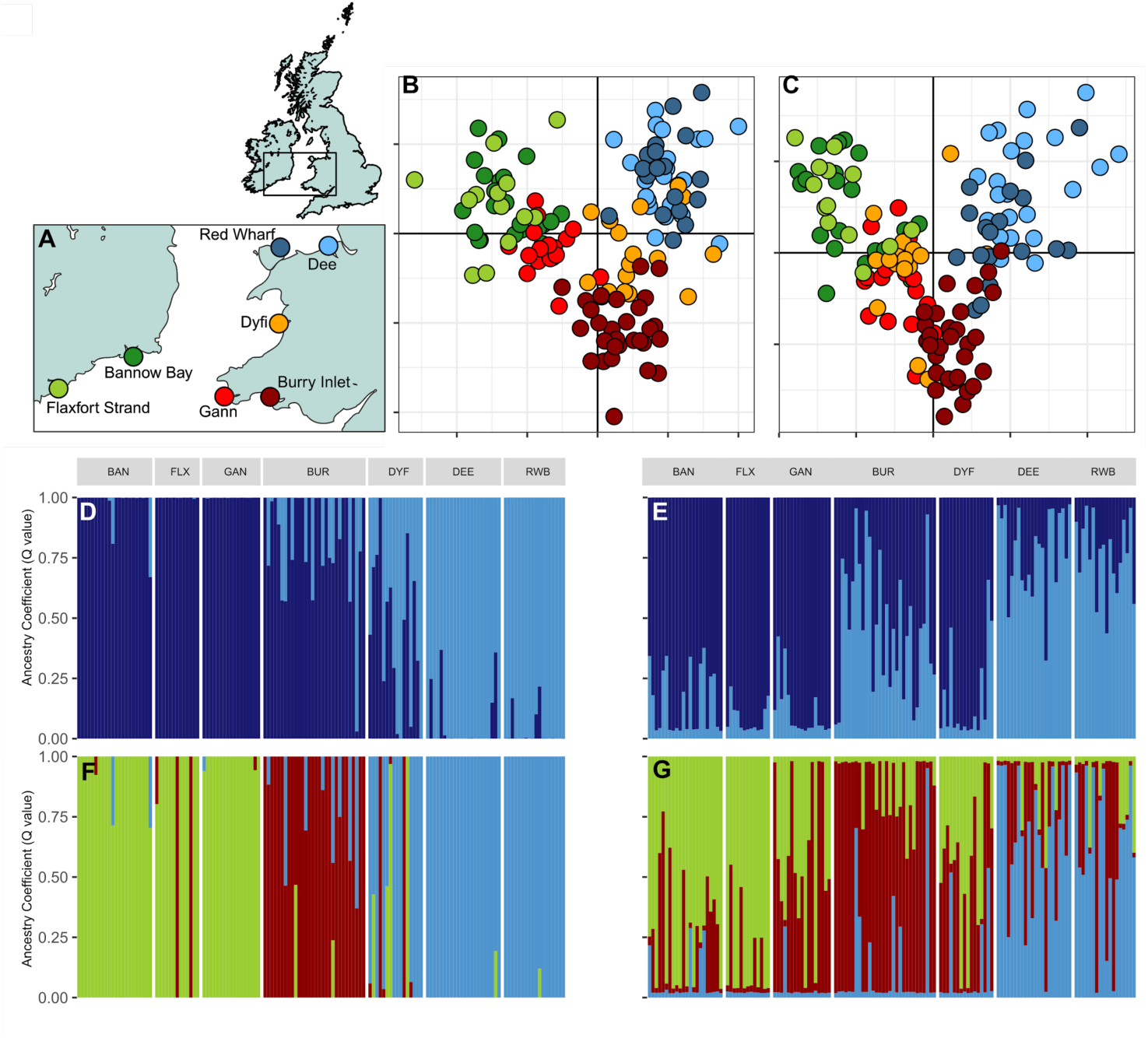
A) Sampling locations; B) DAPC analysis using the neutral dataset; C) DAPC analysis using the 14 outliers; below, the barplots generated by fastSTRUCTURE from the neutral (D and F) and outlier (E and G) datasets, for both K=2 and K=3.

**Table 1:** genetic diversity indices for the two datasets. N, number of individuals remaining after filtering for each location; H_O_, observed heterozygosity, H_E_, expected heterozygosity and F_IS_, the inbreeding coefficient. Values that are significant (95% confidence interval) are in bold.

|  |  | neutral |  |  |  | outlier |  |  |
| --- | --- | --- | --- | --- | --- | --- | --- | --- |
| | | $N$ | $H_O$ | $H_E$ | $F_{IS}$ | $H_O$ | $H_E$ | $F_{IS}$ |
| Bannow Bay | BAN | 22 | 0.133 | 0.150 | **0.087** | 0.269 | 0.272 | 0.006 |
| Gann | GAN | 17 | 0.129 | 0.147 | **0.094** | 0.246 | 0.240 | 0.032 |
| Dee | DEE | 22 | 0.116 | 0.148 | **0.173** | 0.218 | 0.355 | **0.335** |
| Dyfi | DYF | 11 | 0.103 | 0.149 | **0.238** | 0.146 | 0.234 | **0.394** |
| Flaxfort Strand | FLX | 13 | 0.103 | 0.156 | **0.275** | 0.197 | 0.266 | **0.201** |
| Red Wharf | RWB | 18 | 0.118 | 0.144 | **0.141** | 0.286 | 0.337 | **0.179** |
| Burry Inlet | BUR | 30 | 0.109 | 0.150 | **0.230** | 0.173 | 0.218 | 0.159 |

### 2.2 RAD sequencing and bioinformatic analysis

Reduced representation libraries (Baird et al., 2008) were constructed using the restriction enzyme *PstI* (New England Biolabs) for Restriction-Site Associated DNA sequencing (RAD-seq). RAD libraries (each consisting of 24 uniquely barcoded individuals) were produced according to Etter et al. (2011). Each library was quantified using real-time PCR and single-end (100bp target) sequenced on an *Illumina HiSeq2000* at Aberystwyth University, UK. Initial bioinformatic analysis, including quality control, demultiplexing and identification of polymorphisms were performed by *Floragenex* (Eugene, OR; www.floragenex.com), using *Samtools* (Li et al., 2009) and custom scripts, retaining one SNP per tag, with a minimum coverage depth of six for each allele and genotyped in at least 70% of the individuals in the overall dataset. These settings produced an initial dataset of 191 individuals and 4271 single nucleotide polymorphisms (SNPs), which was then further filtered by the authors using the packages *poppr 2.8.1* (Kamvar, Brooks, & Grünwald, 2015; Kamvar et al., 2019; Kamvar, Tabima, & Grünwald, 2014), and *adegenet* (Jombart, 2008; Jombart & Ahmed, 2011; Jombart et al., 2018) in *R* (R Core Team, 2019).

In the second round of filtering, markers with data missing in more than 25% of individuals were discarded from the dataset. A second filtering step removed all loci with *F*_IS_ equal to 1, −1 or NA. Three MAF (minimum allele frequency) filters of 0.05, 0.025 and 0.01 were then applied, generating three datasets. No significant effect of MAF filter upon heterozygosity, global *F*_ST_ and population structure was detected, and so a MAF filter of 0.01 across all sites was applied (Xuereb et al., 2018). This allowed us to maximize the number of markers and hence the information available at low spatial scale, while reducing the bias that might be introduced by retaining low frequency SNPs (Roesti, Salzburger, & Berner, 2012). All the downstream analyses were thus performed on the MAF=0.01 dataset. Markers in linkage disequilibrium (LD) were identified using the function *pair.ia* of the *poppr* package (r^2^ > 0.7). Finally, SNPs that were found to deviate from Hardy-Weinberg Equilibrium (HWE; *P=*0.01) in four out of seven populations were removed (Wyngaarden et al., 2018). The final, filtered dataset included 138 individuals from the seven locations, and 1725 SNPs (Table 1 Supplementary Material).

### 2.3 Neutrality tests and population structure

Genomic markers were tested for neutrality using two complementary methods: *BayeScan* 2.1 (Foll & Gaggiotti, 2008) and the R package *pcadapt.* In order to minimize the detection of false positives, we considered as outliers only those SNPs that were selected by both methods (Coscia, Vogiatzi, Kotoulas, Tsigenopoulos, & Mariani, 2013). *BayeScan* is an *F*_ST_-based, bayesian method (Beaumont & Balding, 2004), while *pcadapt* is based on a Principal Component Analysis (PCA) of individual genotypes and is known to perform particularly well in presence of weak structure, admixture or range expansions (Luu et al. 2017). To set the most appropriate number of clusters (K), we followed the recommendations of the authors and chose the K before the plateau in a scree plot, which displays in decreasing order the percentage of variance explained by each principal component (PC). Q-values were used to account for false discovery rate and SNPs were considered as significant outliers at alpha values ≤0.05. Genetic diversity was estimated on three datasets: overall, neutral and outliers, in order to disentangle the role of demographic processes vs selection. Heterozygosity (expected and observed) was estimated with the R package *hierfstat* (Goudet, 2005; Goudet & Jombart, 2015), while population pairwise *F*_ST_ (Weir & Cockerham, 1984) and relative 95% confidence interval (1000 bootstraps) was estimated with the R package *assigner* (Gosselin, 2019).

Individual-based population structure was assessed using two different approaches: discriminant analysis of principal components (DAPC) as implemented in *adegenet* (Jombart, 2008; Jombart & Ahmed, 2011; Jombart et al., 2018) and *fastSTRUCTURE,* run with a simple prior (Raj, Stephens, & Pritchard, 2014), with the number of k clusters that best explains the structure in the data chosen using the *chooseK.py* script within the *fastSTRUCTURE* software. The power of the neutral and outlier datasets to discriminate and assign individuals was determined with a genotype accumulation curve (Fig. 2 Supplementary Material), calculated with the function *genotype_curve* of the *poppr* package (Kamvar et al., 2015, 2019, 2014).

**Figure 2:**
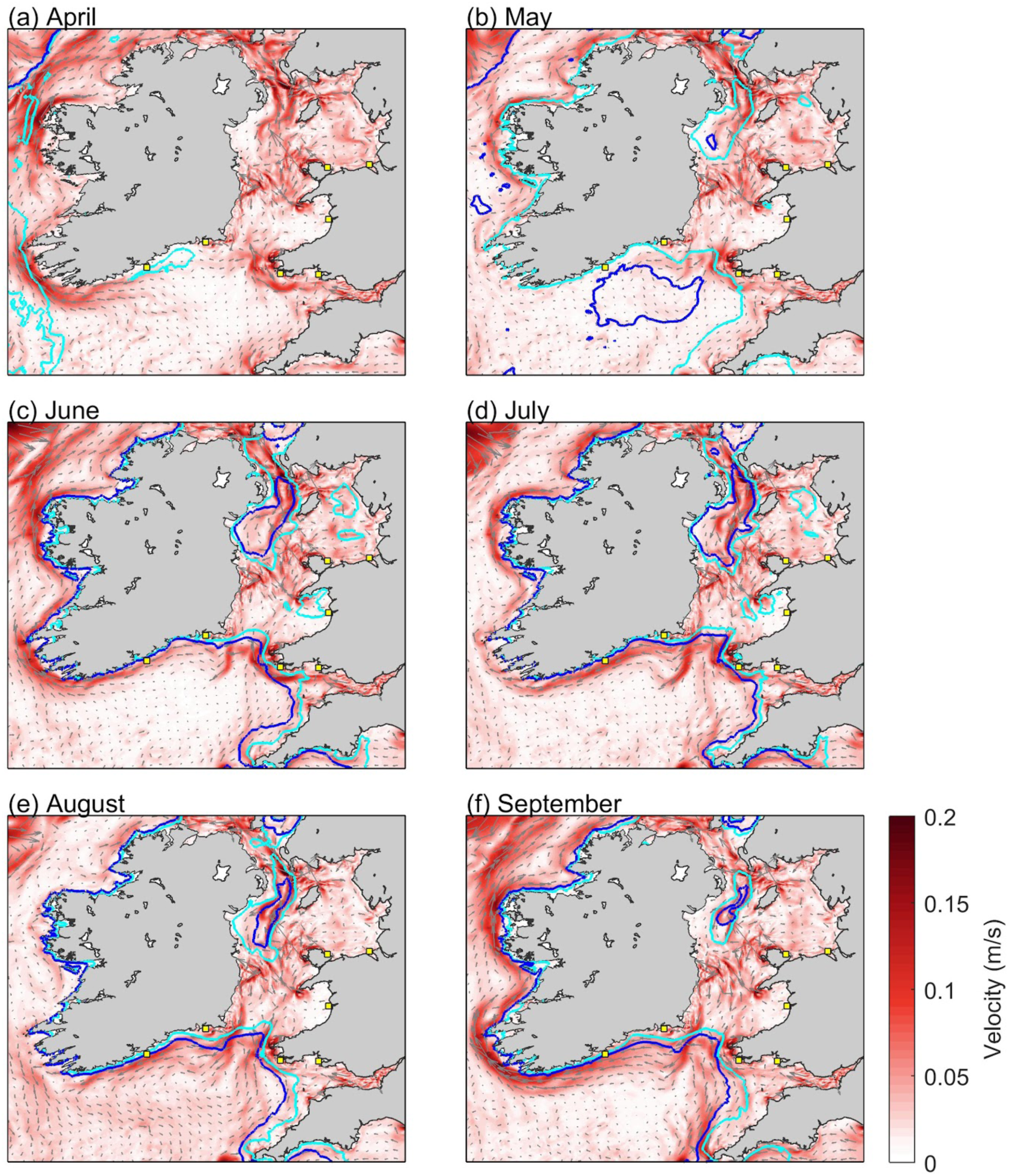
Simulated development of stratification and density-driven currents in the Irish and Celtic Seas, during April-September 2014 (a-f). Light and dark blue contours denote surface-bottom temperature differences of 1°C and 2.5°C, respectively. Each panel shows monthly-averaged and depth-averaged velocity magnitude (red shading) and direction (vectors). Sample locations are indicated with yellow squares.

### 2.4 Hydrodynamic modelling

Hydrodynamic modelling was used to simulate larval transport and predict connectivity between the seven sampled cockle populations, considering natural variability in larval dispersal caused by the timing of spawning relative to the tide, larval duration and seasonal variability of ocean currents. Simulated 3-D flow fields throughout the Irish and Celtic Seas during the cockle spawning/settlement season (April - September) were obtained from the NEMO Atlantic Margin Model (AMM15). The model has a horizontal resolution of 1.5 km and 51 terrain-following layers. It was developed to resolve key dynamical features of the European north-west shelf including the influence of shelf-break dynamics. Bathymetry was provided from EMODnet (EMODnet Portal, September 2015 release) and bottom friction was controlled through a log layer with a non-linear drag coefficient set at 0.0025. Ocean boundary conditions were taken from the Global Seasonal Forecast System (GLOSEA), which includes assimilation of both satellite and in-situ observations. Atmospheric forcing was driven by the European Centre for Medium-Range Weather Forecasts (ECMWF) atmospheric reanalysis product, ERA-Interim (Dee et al., 2011), while river inputs were based predominantly on daily climatology of gauge data. See Graham et al. (2018) for further details of the model development and validation. The simulated flows from AMM15 were used to drive a Particle Tracking Model (PTM) developed to simulate cockle larval transport from each sampled population. In particular, 3-D velocities during April-Septemberwere output at hourly-averaged temporal resolution.

### 2.5 Particle Tracking Model (PTM)

The PTM simulates the lagrangian movement of individual particles in space and time, based on oceanographic dispersion and individual particle behaviour. The PTM was programmed in Python and all simulations were run in a parellised framework on high-performance computing. Hourly-averaged outputs of 3-D velocities from the ocean model was deemed practical in terms of data storage and PTM computational efficiency, whilst minimising uncertainties in particle dispersal generated from interpolating model output.

A previous sensitivity study by Robins et al. (2013) demonstrated that cohorts of 10,000 particles released from a range of point sources within the Irish Sea were sufficient to statistically represent dispersal and connectivity patterns. Accordingly, cohorts of 750 particles were released from sites 1-7 (Fig. 3) each day (at 12:00) over the initial 16 days of April 2014 (i.e. a total 12,000 particles per site covering a spring-neap tidal cycle). This procedure was repeated from April to September 2014. The dispersal of each particle was tracked for 40 days pelagic larval duration (PLD), reflecting the observed PLD of cockle larvae (Malham et al. 2012). Particles were neutrally buoyant and able to disperse throughout the 3-D flow field. Connectivity between populations was determined from particle trajectories during days 30-40; particles that travelled within 20 km of a settlement site were assumed to have settled there. The 20 km threshold was based on the average tidal excursion for the Irish Sea (Robins et al. 2013), although a range of thresholds are discussed in the Results section.

**Figure 3:**
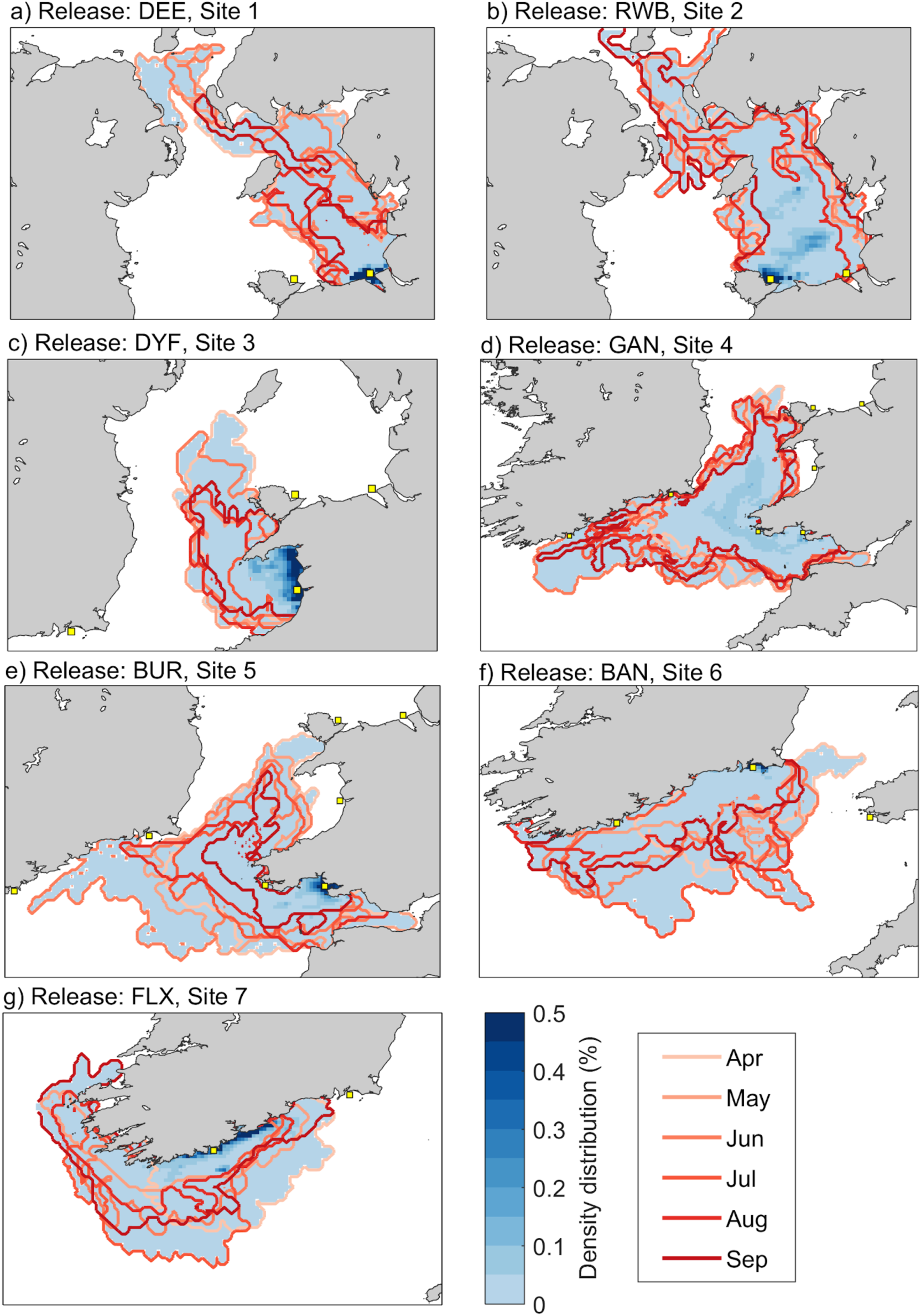
Probability density distribution maps showing simulated dispersal probability from release sites 1-7 (yellow squares). Each panel (blue colour scale in a-g) shows dispersal range for 11,520,000 particles (12,000 each month × 6 months × 10 settlement days). The superimposed contour lines show the maximum spread per month, hence an indication of seasonal dispersal variability.

### 2.6 Modelling uncertainty

The ocean model AMM15 was run in free mode without data-assimilation. When its results were compared with a simulation with data-assimilation for a different year to this study, the inherent biases were small and are not expected to change the advection along the tidal mixing fronts and hence larval transport. For the larval dispersal scenarios considered here, the simulated variability in dispersal distance after 40 days, due to the release day (particles were released daily for 16 days), was of the same order as the variability encountered within each daily cohort of particles. This result demonstrates the importance of ‘trickle-spawning’ larvae over a full lunar tidal cycle to capture dispersal variability.

In the absence of strong evidence for *C. edule* larval swimming behaviour, the effects of larval behaviour were not incorporated into the particle tracking model. The general agreement in population structure between our genomics and modelling analyses could be interpreted as evidence for a larval transport mechanism that is controlled by ocean dispersion rather than swimming behaviour. In effect, this result suggests that the ocean currents are much stronger than larval swimming speeds.

In this work, one year of larval dispersal was simulated. Due to interannual variations in weather patterns the timing of the onset of seasonal stratification and its strength can vary between years, variability in dispersal due to atmospheric conditions over one season has been accounted for in our study. As 2014 was an ‘average’ year with respect to the long-term current and stratification variability, we are confident that this year is representative of the mean transport pathways in the Irish and Celtic Sea region. However, future work could investigate the influence of interannual variations in ocean climate on larval dispersal. since, in the aftermath of severe storm events, ocean circulation patterns have been shown to reverse in UK shelf-seas (Stanev et al., 2019). Although such storms are rare, having only occurred around the UK four times in the last 40 years, changes in ocean circulation during spawning cycles due to rare and severe storms create additional uncertainty in larval trajectories, potentially leading to the establishment of new communities which have no clear connection at other times.

Despite the relatively fine spatial resolution of our model (1.5 km with 51 vertical layers), some coastal currents will inevitably be poorly resolved, adding a potential source of uncertainty for larval transport in the coastal zone. Coastal modelling requires an unstructured grid mapped onto a high-resolution coastal bathymetry and including wetting/drying capabilities within the model and also wave-induced currents. This is a considerable task in terms of validation and application to particle tracking, but should be considered for further studies.

### 2.7 Spatial Eigenfunction Analysis (SEA) and Environmental Association (EA) analysis

To test for association between the oceanographic environment and the cockle genetic structure, a Spatial Eigenfunction Analysis (SEA) (Dray et al., 2012) was performed. This approach was implemented through the *vegan* and *adespatial* packages (Dray et al., 2019; Oksanen et al., 2017) in R. SEA was used to estimate the influence of geographic distance between samples as well as the influence of modelled connectivity on genomic variation. Geographic distance was represented as distance-based Moran’s eigenvector maps or dbMEM (Stéphane Dray, Legendre, & Peres-Neto, 2006). dbMEMs were calculated using the *PCNM* function of the *vegan* package on the Euclidean distances (calculated with the function *dist*) in turn estimated from Cartesian-transformed coordinates using the *geoXY* function in the *SoDA* package (Chambers, 2013). *PCNM* transforms spatial distances in rectangular matrices that are suitable for constrained ordination (Legendre & Gallagher, 2001).

For the Environmental Association analysis (EA), modelled connectivity was represented as asymmetric eigenvector maps or AEM (Blanchet, Legendre, & Borcard, 2008; Blanchet, Legendre, Maranger, Monti, & Pepin, 2011). AEM is a spatial eigenfunction approach specifically developed to model multivariate responses to asymmetric and directional processes such as current-driven larval dispersal (Blanchet et al. 2011). The connectivity probability matrix generated by the biophysical model was translated into a nodes-to-edge matrix which records the presence/absence of connectivity links (through *edges*, 19 here) between *nodes* (the 7 sampling locations). Each edge has an associated ‘weight’, based on the simulated probability of connectivity. AEMs were calculated with the *aem* function in R.

For each sampling location, the simulated monthly-averaged sea surface temperature (SST) and surface-bottom temperature difference (SBTD - the difference between the temperature at the surface and the bottom of the water column) representing ocean stratification, from April to September, were extracted from the ocean model (Figure 1 Supplementary Material). The relative contribution of temperature (SST and SBTD), spatial distribution (dbMEMs) and directional connectivity (AEMs), to genetic variation in both the neutral and the outlier datasets (response variable) was modelled using redundancy analysis (RDA). In particular, the response variable was represented by population-specific minor allele frequencies (MAF) of each SNP calculated in the R package *hierfstat* (function *minorAllele*) (Goudet and Jombart 2015), detrended using the *decostand* function with the *hellinger* transformation available in *vegan* (Oksanen 2018). The most important explanatory variables were chosen by performing the forward selection with 10,000 permutations in *vegan* using the function *ordistep.* Each model’s significance was assessed with an analysis of variance (function *anova* in *vegan*) with 1000 permutations, to finally establish which factors were most correlated with genetic variation. Redundancy analysis (RDA) and partial-RDA (corrected for geographic distance between populations) were performed using the *rda* function in *vegan* on the putatively neutral (1711) and outlier (14) SNP datasets, and parsimonious RDAs were carried out using the variables selected (Borcard, Gillet, & Legendre, 2011).

## 3. RESULTS

### 3.1 Genetic diversity and population structure

After filtering for missing data, *F*_IS_, MAF, HWE and LD, 138 individuals and 1725 SNPs were retained (Table 1 Supplementary Material). *Bayescan* detected 28 SNPs as potential outliers, with a False Discovery Rate of 0.05, whereas *PCAdapt* found 62. Only 14 SNPs overlapped between the two approaches. The downstream analyses were then carried out on two datasets: neutral (1711 SNPs) and outliers (14 SNPs).

For the neutral dataset, expected and observed heterozygosity (*H*_E_ and *H*_O_) were similar across locations, ranging between 0.148-0.157 and 0.103-0.134, respectively (Table 1). Pairwise *F*_ST_ (Fig. 3 and 4 Supplementary) ranged between 0 (RWB-DEE) and 0.0289 (GAN-DEE). *fastSTRUCTURE’*s *choosek.py* detected three clusters (k=3; Fig. 1D-G), while *DAPC*’s *find.clust* found a maximum of five clusters (k=5; Fig. 1B-C) based on the BIC scores. For the putative outlier dataset, *F*_ST_ values ranged between 0 (DYF-GAN) and 0.38 (BUR-FLX), but *choosek.py* and *DAPC*’s *find.clust* indicated that one cluster explained the structure in the data (barplots represented in Fig. 1E and 1G for comparison). All population-level neutral *F*_IS_ values were significant (95% confidence interval) and positive, ranging between 0.08 and 0.27 (Table 1). For the outlier dataset, *H*_E_ and *H*_O_ varied respectively between 0.21-0.35 and 0.14-0.28 and *F*_IS_ values were positive and significant for four populations: DEE, RWB, DYF and FLX.

**Figure 4:**
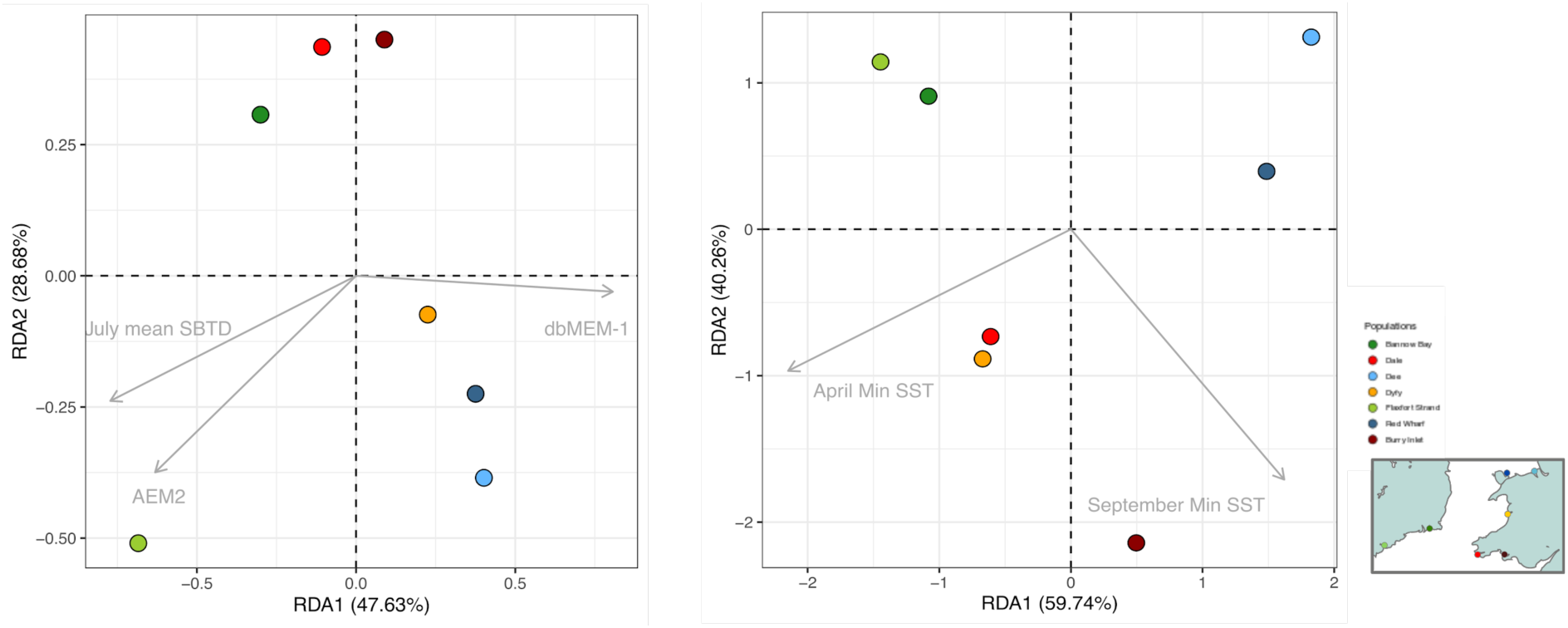
Redundancy analysis (parsimonious RDA) performed for the (A) neutral and (B) outlier datasets. Each circle is a sampling location, and each arrow is an environmental variable that significantly drives the observed population structure.

Genotype accumulation curves (Fig. 2 Supplementary Material) for both datasets show that a plateau is reached and variance decreased at 20 neutral and 12 outliers loci respectively, indicating that the datasets have sufficient power to distinguish between individuals (Arnaud-Haond, Duarte, Alberto, & Serrão, 2007).

### 3.2 Seasonal circulation and larval dispersal

The typical circulation structure within the Irish/Celtic Seas has, in our study, led to distinct patterns of larval transport and population connectivity, but with spatial variability over seasonal timescales. Whilst the tidal currents in the Irish Sea are characteristically large (e.g. peak flows exceed 1 m/s throughout much of the basin; Robins et al. 2015) and have the potential to advect larvae tens of kilometers during each tidal cycle, these currents are oscillatory, and resulting in small net flows and minimal dispersal over several weeks (Robins et al. 2013). However, the development of thermal stratification and ocean fronts during summer months generates density-driven currents along frontal boundaries (Horsburgh, Hill, & Brown, 1998; Simpson & Hunter, 1974). Although much weaker than tidal currents, these density-driven currents are persistent over time and act as key dispersal pathways for larvae. For example, the Celtic Sea front may facilitate connectivity of cockle populations from South Wales across to Ireland (Coscia et al. 2013).

The AMM15 simulation shows the Celtic and Irish Seas during April 2014 to be vertically well mixed with net tidal flows following well-known patterns due to tidal interactions with morphology (Pingree & Griffiths, 1979) (Fig. 2A). These net flows were generally less than 0.1 m/s and directed south/southwest and north/northeast from a parting zone (amphidromic point) south of Dublin. During May-September, the simulation shows the development of thermal stratification leading to density-driven currents (Fig. 2B-F). The Celtic Sea and the western Irish Sea stratified generating persistent east-to-west currents along the Celtic Sea front and an anti-clockwise gyre in the western Irish Sea. For the Celtic Sea front, depth-averaged flows were ∼0.15 m/s but strongest at the thermocline at ∼30m depth. These density-driven flows persisted but weakened during August and September.

The degree of exposure of the spawning grounds influenced the population’s ability to connect with other sites (Figs. 3 and 5 Supplementary). This is shown by comparing the simulated particle dispersal from DEE (Site 1) with RWB (Site 2) on the north Wales coast. The majority of particles dispersing from DEE remained <30 km from the release site, resulting in high proportion of self-recruitment (17.4±0.7%), with only small proportions of particles dispersed north to the North Channel and the Scottish coast (Figs. 3A and 5 Supplementary). In contrast, particles from RWB were exposed to stronger currents and the site experienced a lower degree of self-recruitment (13.6±3.5%) with ‘hot-spots’ of higher density particles (up to 0.5%) advected ∼100 km northwards (Figs. 3B).

Our simulations suggest a high degree of isolation at DYF (Site 3) (Figs. 3C and 5 Supplementary), with the majority of particles retained within Cardigan Bay. Due to the weak flows within Cardigan Bay, simulated particles from other sites did not reach DYF within the 40-day larval stage. Simulated particles from GAN (Site 4) and BUR (Site 5) were capable of dispersing readily between one another (up to 1% connectivity) and also westwards to the Irish populations, particularly from the more exposed GAN (up to 0.5% connectivity) (Figs. 3D-E and 5 Supplementary). Connectivity with DEE was possible but unlikely (<0.01%). It was clear from the simulations that seasonal patterns of dispersal are expected, e.g. from BAN (Site 6) and FLX (Site 7) (Figs. 3F-G and 5 Supplementary). Simulated dispersal from these populations was controlled by the persistent westward residuals around southern Ireland, with the particles generally travelling further west as the residuals strengthened during summer months. For the BAN population, only during April when the Celtic front had not formed, were particles able to disperse northwards into the Irish Sea.

### 3.5 Environmental Association analysis

The *ordistep* function found that three variables were linked to the population structure found at neutral loci: July SBTD explained 10% of the genetic variance (*P*=0.002, adj*R*^2^=0.10), dbMEM-1 7% (*P*=0.000, adj*R*^2^=0.07) and AEM2 4 %(*P*=0.050, adj*R*^2^=0.04). Partial RDAs on neutral data using these three variables were not significant.

For the outlier dataset, two environmental factors were found to be highly significant by *ordistep*, explaining 71% of the genetic variance: April min SST (*P*=0.025, adj*R*^2^=0.71) and September min SST (*P*=0.004, adj*R*^2^=0.71). These SST values were the lowest daily mean SST in each month. The global parsimonious RDA (Fig. 4A) was overall non-significant when including the three selected factors at once (*P*=0.11), although it was significant when including two at a time: dbMEM-1+July mean SBTD (*P*=0.008), AEM2 + dbMEM-1 (*P*=0.022). Partial RDAs were not significant. The parsimonious RDA run on outliers was globally highly significant (*P*=0.002; Fig. 4B).

For the neutral RDA, the first axis explained almost 48% of the total variance, and was mainly driven by dbMEM-1 geographic distance between sites. This axis separates the northern sites of Red Wharf Bay, Dee Estuary and Dyfi Estuary from the southern sites of Flaxfort Strand, Bannow Bay, Gann Estuary and Burry Inlet, describing a north-south divide that is not connected by oceanic currents. The second axis (RDA2) was influenced predominantly by an asymmetrical eigenvector representing modelled connectivity (AEM2) and July sea surface-bottom temperature difference (July mean SBTD), a key determinant of ocean stratification. On this axis, the southernmost population Flaxfort Strand (light green) is strongly related to AEM2, the asymmetrical eigenvector that represents larval dispersal (Fig. 4A). Redundancy analysis carried out on the outliers dataset revealed patterns of local adaptation, with cool sea surface temperatures being the most significant driver (Fig. 4B).

## 4. Discussion

This is the first study to use genomic markers and biophysical larval transport modelling to investigate the drivers of connectivity in a commercially important shellfish in north-western Europe. This research was conducted in the Irish and Celtic Seas where *C. edule* forms valuable shellfisheries for both Ireland and the United Kingdom. A panel of 1725 SNP markers was analysed in relation to important environmental variables such as temperature and oceanographic currents, factors that have been shown to be drivers of population structure in bivalves (Araneda et al., 2016; Gormley et al., 2015; Bernatchez et al., 2019; Lehnert et al., 2019; Xuereb et al., 2018). Using genomic markers, three main genetic groups were identified, which can be considered stocks or management units. Overall, the genetic results fit well with the predictions of the larval transport model, providing a level of empirical validation for both the simulated hydrodynamics and larval behavioural traits. Environmental association analysis revealed that neutral genetic structure was strongly linked to geographical distance between sites and to the strength and direction of the ocean currents acting as corridors for larval dispersal, whereas colder periods (cold SSTs) were identified as the drivers of local adaptation.

### 4.1 Connectivity, fine-scale population structure and local adaptation

Neutral genetic diversity was very low across the study area, and compared with previous studies on marine bivalve population genomics (Bernatchez et al., 2019, Lehnert et al., 2019). Populations of cockles in the Irish Sea have been under pressure for at least two decades, with mass mortality events and declines due to overexploitation leading to strict management of most beds across the UK (Woolmer, 2013). In addition, variance in reproductive success is known to occur in bivalves (Hedgecock & Pudovkin, 2011). These events could be responsible for the loss of genetic diversity, as already observed in several marine organisms (Pinsky & Palumbi, 2014). On the other hand, heterozygote deficiency (as indicated by positive *F*_IS_ in all sampling locations - Table 1) is well known to occur in marine bivalves (Gaffney, 1994), and has already been detected in these populations of common cockles using microsatellites (Coscia et al. 2013). Furthermore, cockles have been shown to undergo boom and bust years (Morgan, O’ Riordan, & Culloty, 2013) with the dispersal of cockle larvae and recruitment altering between years and with external parameters such as temperature (Miller, Versace, Matthews, Montgomery, & Bowie, 2013; Morgan et al., 2013).

Given the fine spatial scale and the reproductive biology of the study organism (broadcast spawner with a long pelagic larval phase), a lack of, or weak population structure was expected. Previous studies in the same geographic area identified a lack of genetic structure in shellfish species using microsatellite markers (Coscia et al., 2013; Watson et al., 2016; McKeown et al., 2019). In this study, three main genetically distinct units were identified with the neutral marker set. The first group includes the north Wales populations of Red Wharf Bay and Dee Estuary, the second includes Bannow Bay and Flaxfort Strand on the south coast of Ireland and the Gann Estuary in the south-west of Wales, and the final group contains only the Burry Inlet on the south coast of Wales. To our knowledge, this is the first time such a pattern has been detected in *C. edule*, or in any other shellfish resource in the Irish Sea.

Geographic proximity certainly favours gene flow, but oceanographic currents aiding larval transport are also major drivers of population structure (Barbut et al., 2019). For example, Red Wharf Bay and the Dee Estuary appear to be genetically homogeneous, due to high levels of gene flow, but these populations are very distinct from those further south (150-350 km away). This concurs with the model’s prediction that prevailing currents will mostly disperse larvae from the north coast of Wales northwards. The high levels of gene flow detected between the Gann Estuary on Wales’ southwest coast and sites on the southeast coast of Ireland also correspond with the model’s prediction of high levels of westward dispersal along the Celtic Sea front (Coscia et al. 2013).

Of particular interest is the genetic make-up of the population in the Burry Inlet. Here, cockles have experienced recurrent mass mortality events for 15 years (https://marinescience.blog.gov.uk/2015/08/14/unusual-cockle-mortalities-burry-inlet/). The relatively high level of genetic differentiation between the neighbouring Burry Inlet and Gann estuary populations indicates that gene flow between these populations is low, seemingly contradicting the larval transport model’s prediction of high connectivity. This result suggests that the Burry Inlet population may able to persist through self-recruitment, rather than forming a sink population depending upon immigration from healthier populations elsewhere, and could explain the low genetic diversity and high levels of inbreeding detected there. However, the model showed that larval dispersal is possible from the Gann to Burry estuaries, although we assumed a relatively large settlement zone that extended beyond the mouth of the Burry Inlet (Fig. 6 Supplementary). Furthermore, it must be acknowledged that the Gann estuary cockle population is far smaller than that of the Burry Inlet so larvae dispersing from the Gann into the Burry may be swamped by self-recruitment, or might not survive due to local selection against them. Additionally, the southern coast of Wales contains other large cockle populations, such as the Three Rivers fishery mid-way between the Gann and Burry. These populations were not sampled/modelled in this study but may provide a nearer and greater source of larvae for the Burry Inlet than does the Gann Estuary.

The Spatial Environmental Association analysis identified several environmental factors highly associated with the population structure observed at neutral and outlier genetic markers. This is a strong statistical approach, which has already been successfully employed to study the influence of the environment on the genetic structure of commercially important bivalves, such as the eastern oyster (*Crassostrea virginica*; Bernatchez et al. 2019) and Atlantic deep-sea scallop (*Placopecten magellanicus*; Lehnert et al. 2019). In *C. edule,* neutral genetic structure is strongly dependent on geographic distance between sites (dbMEM-1), indicating that isolation by distance plays an important role in shaping the observed genetic structure in this species, despite its long pelagic larval duration. Nevertheless, it is the interplay between isolation by distance, peak temperature-driven currents (July SBTD) and modelled connectivity (AEM2) that shapes neutral population structure. The summer stratification which strengthens the Celtic Sea front current, directed from south Wales to Ireland (Simpson and Hunter 1974), plays a major role in connecting cockle populations between the south of Wales and Ireland, while separating them from populations further north in the Irish Sea. Considering the 14 outlier loci, the minimum sea surface temperatures recorded in April and September explained the genetic variance observed within this dataset. In particular, the Burry Inlet was strongly associated with the SSTs in September, which are warmer compared to the ones recorded at other locations for the same time (Figure 1 Supplementary). If cockles in the Burry Inlet are indeed adapted to warmer sea surface temperatures than populations at sites further west or north (and especially compared to those at Gann), this may explain the maintenance of genetic differentiation between the Burry Inlet and the Gann Estuary despite their spatial proximity and the potential for high connectivity predicted by the model (Fig 3 and Fig. 5 Supplementary Material). Larvae arriving adrift from Gann to the Burry may not be able to survive post-recruitment given selective pressure against them by higher temperatures.

### 4.2 Implications for management

The results of the biophysical modelling are likely to have the greatest relevance to fishery management in terms of the potential seasonal variability in larval supply to cockle grounds. This study demonstrates the existence of three distinct units of cockles using both genomic tools and larval dispersal modelling. As with other recent studies (Lal, Southgate, Jerry, Bosserelle, & Zenger, 2017) these findings have important implications for fishery management (Coscia et al. 2013; Miller et al. 2013) and how fisheries management can be reconciled with conservation and other activities.

Given the incidence of recurrent mass mortality events at the Burry Inlet (Callaway et al. 2013), the genetic isolation of this cockle fishery implied by the results of this study should be investigated further. This could be achieved by expanding the sampling coverage of the Burry Inlet to multiple sites and years and assessing its connectivity to other nearby cockle beds that have not been included in this study. Future investigations should be aimed at clarifying the role of local adaptation into the fine-scale population dynamics of the common cockle in this area, in order to improve management, while also assessing the role played by diseases and infections. The results from this study highlight the importance of the use of genomic and hydrodynamic data in assessing population structure and connectivity in an exploited and commercially important marine species and may aid in current and long-term management regimes of this species (Lal et al. 2016; Bernatchez et al. 2017).

## Data availability

The data that support the findings of this study are openly available in Dryad at http://doi.org/[doi], reference number [reference number].

## Acknowledgements

The genomic data was generated thanks to an internal grant awarded to IC by Aberystwyth University. The authors also wish to acknowledge the support of the Interreg Ireland-Wales Programme ISPP (http://ispp.bangor.ac.uk/) and Bluefish (bluefishproject.com) projects, the Interreg Atlantic Area Programme Cockles project (https://cockles-project.eu/), and the SUSFISH project. Oceanographic data was provided through the SEACAMS project (www.seacams.ac.uk), funded by the Welsh Government, the Higher Education Funding Council for Wales, the Welsh European Funding Office, and the European Regional Development Fund Convergence Programme. The model simulations were conducted on the Supercomputing-Wales high performance computing(www.supercomputing.wales) system (a collaboration between Welsh universities and the Welsh Government), supported by Ade Fewings and Aaron Owen. The authors are grateful to all the people that have helped with sampling, Mandi Knott from the North Western Inshore Fisheries and Conservation Authority and Emer Morgan from University College Cork, as well as in the lab (Matt Hegarty and Elizabeth Harding), and the colleagues Niall McKeown and Hayley V. Watson for continuous help and support. We also are extremely grateful to Paulino Martinez for constructive comments on the manuscript.

## SUPPLEMENTARY MATERIAL

**Figure 1 Supplementary:**
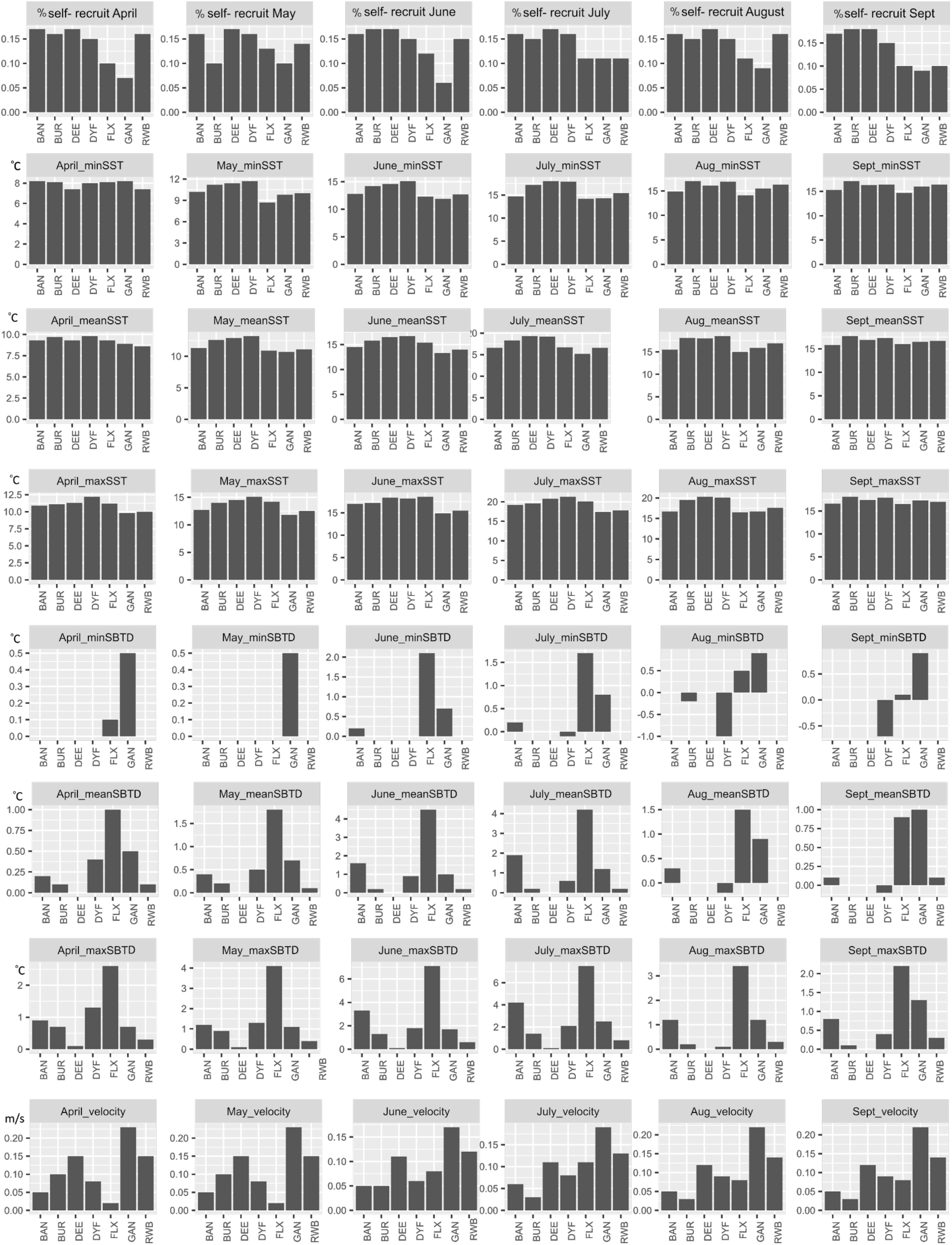
Environmental parameters for each location, and each month used in the environmental association analysis.

**Figure 2 Supplementary.**
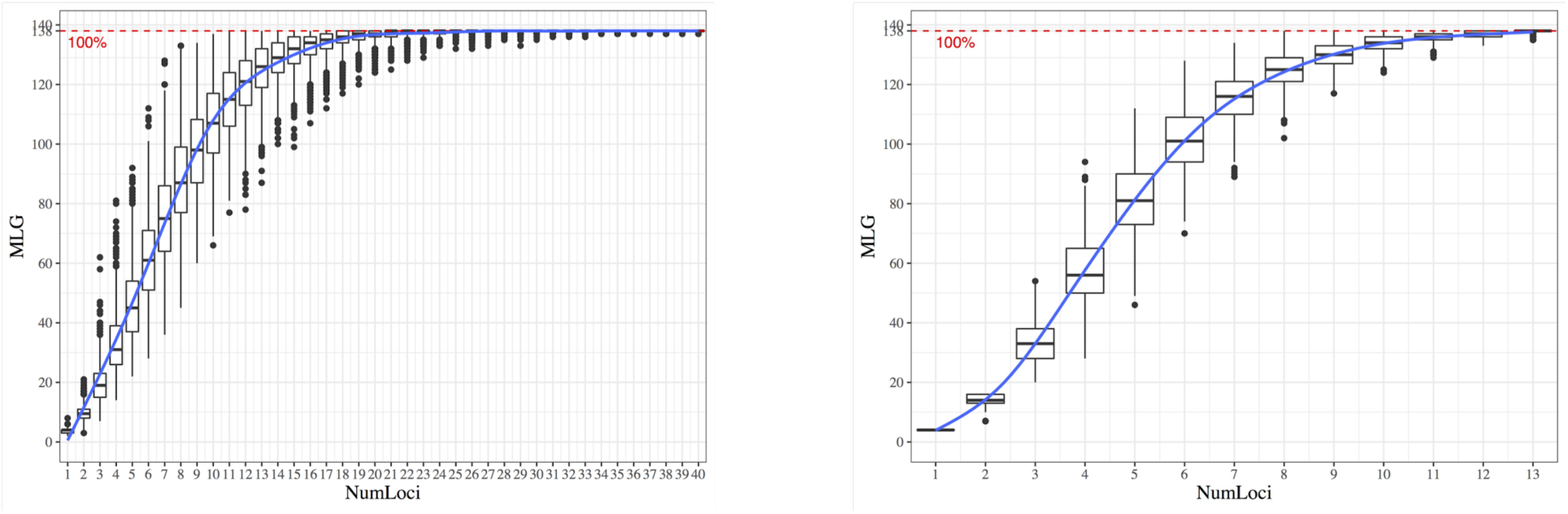
Genotype accumulation curves for neutral (A) and outlier (B) datasets. On the Y axis, the number of multilocus genotypes and on the X axis, the number of loci.

**Figure 3 Supplementary:**
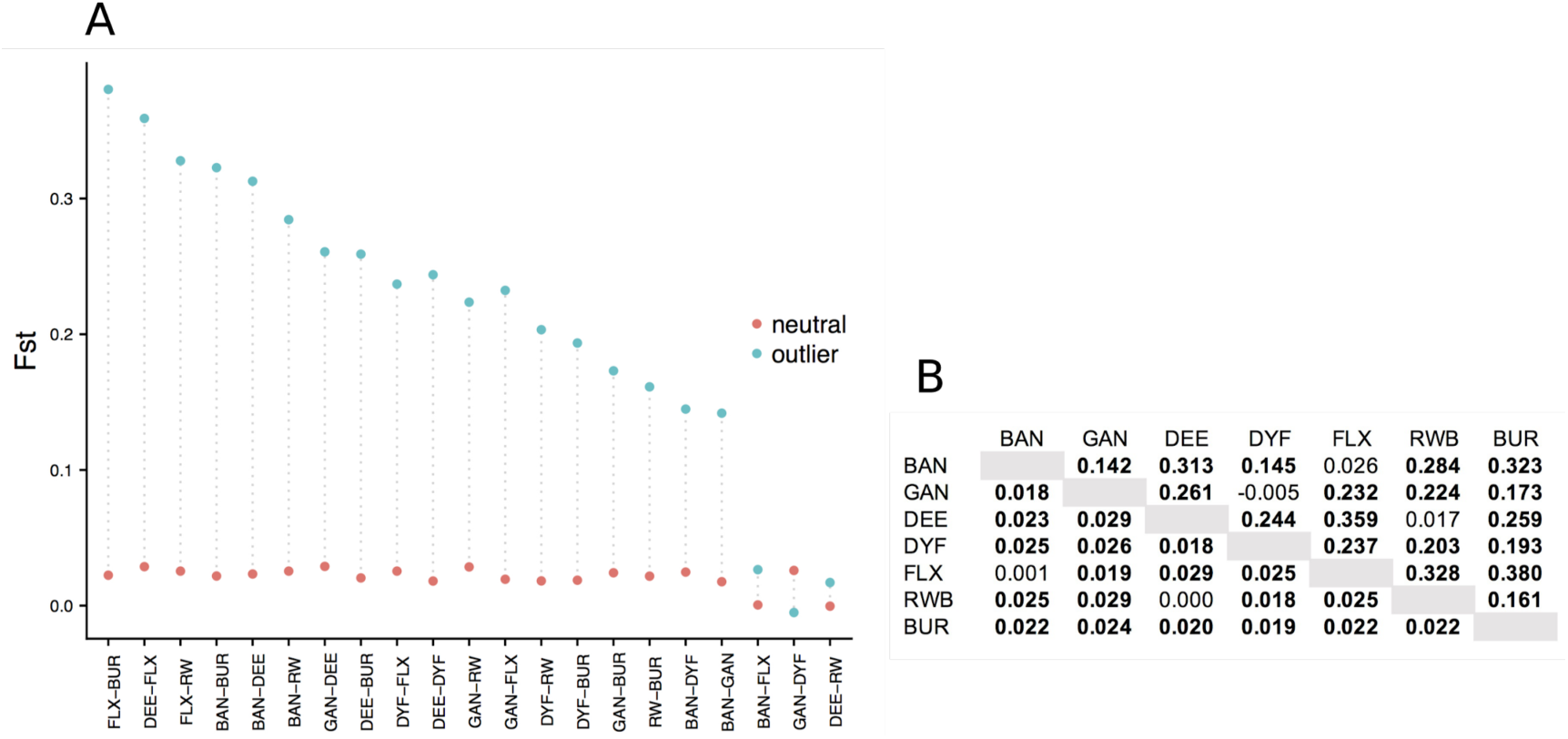
Pairwise Fst calculated for neutral and outlier datasets. The graph (A) shows which pairs have the highest difference between the two. In the table (B) the relative values for neutral (lower diagonal) and outliers (upper diagonal). In bold, values that are significant (95% confidence interval).

**Figure 4 Supplementary.**
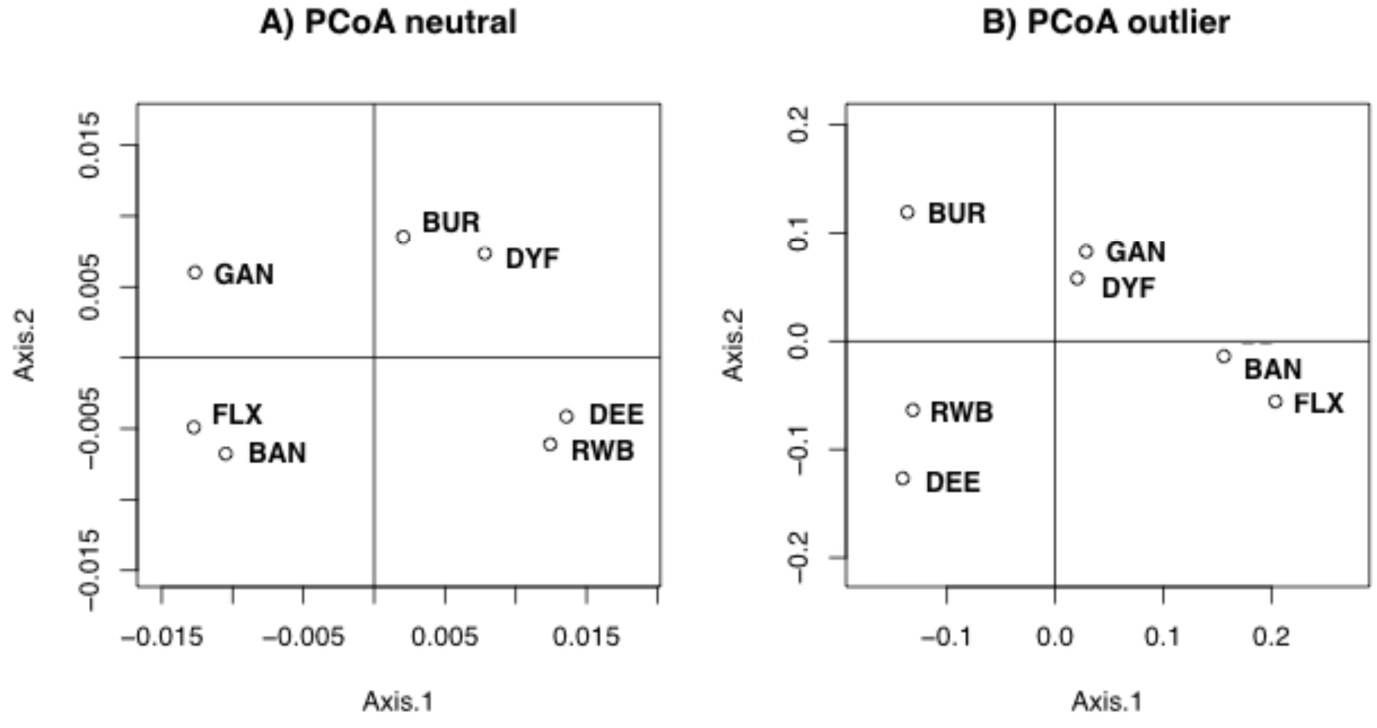
Principal Coordinates Analysis (MDS plots) based on *F*_ST_

**Figure 5 Supplementary:**
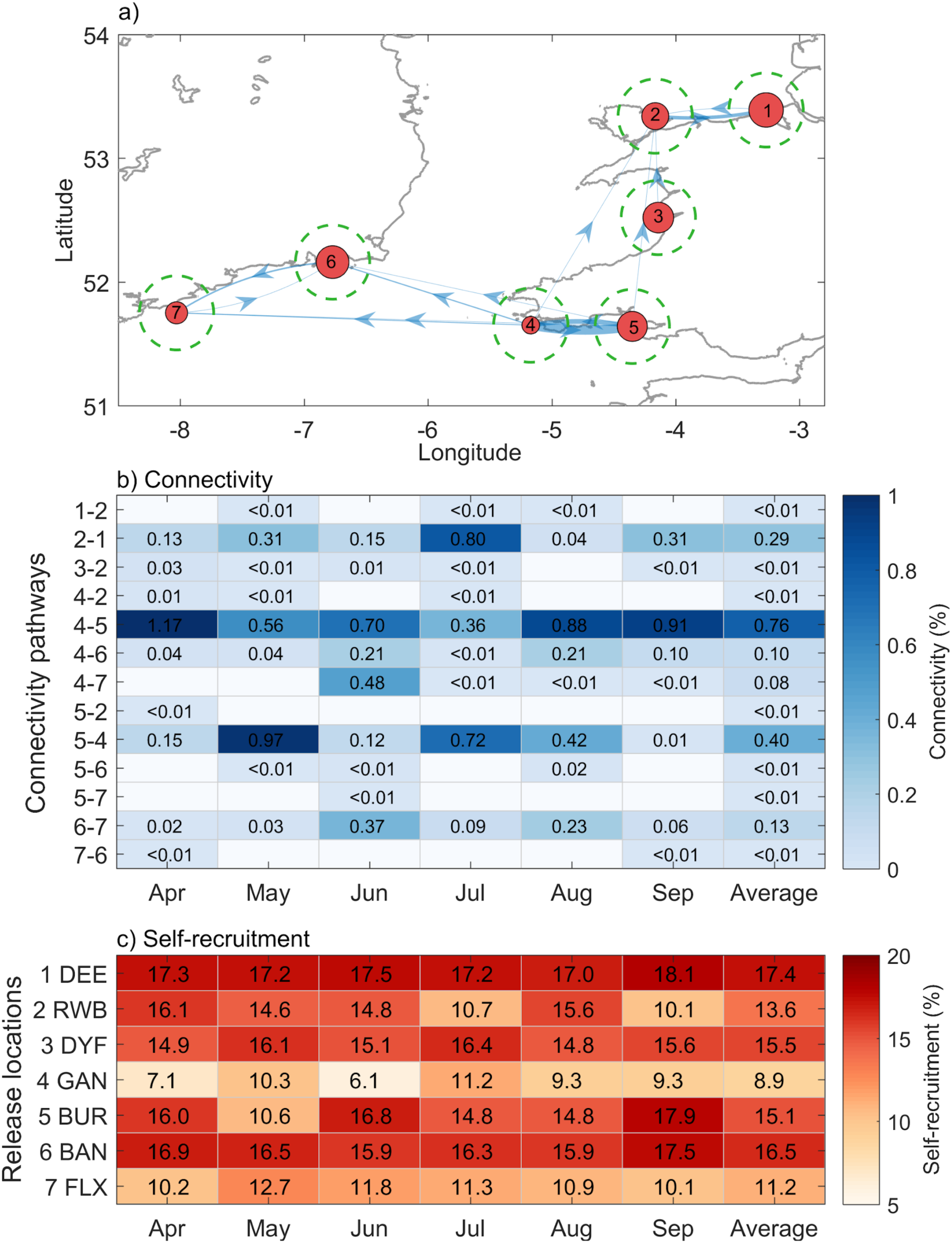
Simulated connectivity between the seven sampled cockle populations. Seasonally-averaged (2014) connectivity networks are shown geographically in (a). The thickness of the pathways in (a) corresponds to the average values in (b). Self-recruitment in (a) is denoted by the size of the red circles which correspond to the average values in (c). The dashed green circles show the settlement radius used to calculate connectivity. Seasonal variability (Apr.-Sep.) is shown for connectivity (b) and self-recruitment (c).

**Figure 6 Supplementary:**
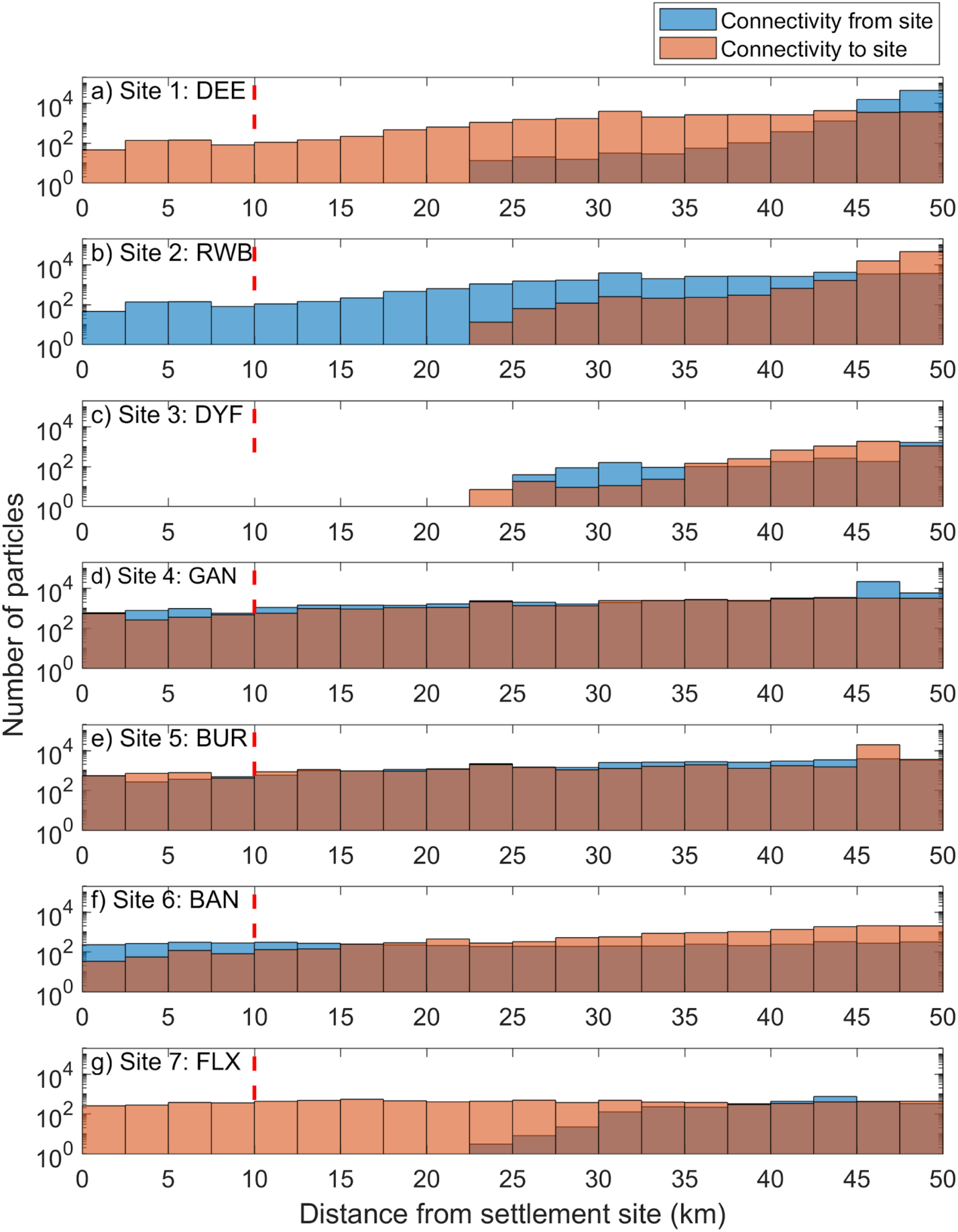
Histogram plots showing connectivity as a function of distance. For each particle trajectory, the minimum distance to a settlement site during days 30-40 is presented. Blue bars signify the connectivity potential of source sites 1-7 (a-g). Red bars signify the connectivity potential of settlement sites 1-7 (a-g). For example, (a) shows that particles released from DEE simulated no connectivity within 20 km of a settlement site, whereas particles from elsewhere did connect with DEE. The dashed red line denotes our threshold radius for connectivity: 10 km away from each settlement site.

**Table 1 Supplementary.** Filtering steps. *#individuals*, remaining number of individuals after each filtering step; *#SNPs* remaining number of SNPs

|  | # individuals | # SNPs |
| --- | --- | --- |
| **Raw dataset** | 191 | 4271 |
| **Missing data** | 159 | 4271 |
| *F<sub>IS</sub>* | 138 | 3488 |
| **MAF 1%** | 138 | 1864 |
| **Linkage Disequilibrium** | 138 | 1775 |
| **Hardy-Weinberg eq.** | 138 | 1725 |

